# The Effect of Human Presence and Activity Type on Innovative Problem-Solving of Urban Eurasian Red Squirrels

**DOI:** 10.1101/2024.11.20.624604

**Authors:** Pizza Ka Yee Chow, Olli J Loukola, Cwyn Solvi

## Abstract

1. Humans impact wildlife positively and negatively, and increasing evidence shows that humans potentially play a major role in shaping cognition in wild animals. Recent evidence suggests, in particular, that the ability to innovate (an important cognitive ability for urban wildlife) can be affected by humans. However, the specific anthropogenic factors associated with shaping animal cognitive performance remain unclear.
2. Here, across 15 urban areas in Oulu, Finland, we recorded 64 wild Eurasian red squirrels’ interactions with a novel puzzle box (innovative problem-solving) to obtain a highly preferred food, hazelnuts.
3. We then examined how different levels of human presence nearby, types of activity (walking, dog walking, cycling, and playground activities), and distance to the nearest footpaths influenced squirrels’ innovative problem-solving ability – measured as solving success at the population level and solving latency at the individual level.
4. At the population level, higher mean human presence nearby significantly decreased the proportion of squirrels that successfully solved the novel food-extraction problem (“innovators”), and greater human presence nearby decreased first-visit solving latency at individual level. Except for cycling, all types of human activities (walking, dog walking, and playground activity) significantly decreased first-visit success rate. Dog walking and playground activity had a greater negative impact on success rate at the population level than walking. Dog walking was the only outstanding factor affecting first-success latency. We also detected interaction effects between the types of human activities and nearest footpath on overall success rate of all innovators.
5. These results provide evidence for the negative effect of specific human-related factors on squirrels’ innovative problem-solving abilities. These factors may also exert selective pressure, potentially shaping the evolution of wildlife cognition.

## Introduction

Urban environments have been shown to affect many aspects of wildlife, including physiology (Patankar et al., 2021), morphology (Biard et al., 2017; Evans et al., 2009), and behaviours (Candolin & Wong, 2012; Caspi et al., 2022; Lowry et al., 2012; Ritzel & Gallo, 2020; Sih, 2013; Sih et al., 2011). A recent growing body of evidence suggests that the impacts of urbanisation can extend to cognition (Chow et al., 2024; Chow et al., 2021; Ducatez et al., 2015; Papp et al., 2015). Cognition manifests through behaviours, and cognition can be defined as an individual acquiring, processing, storing, utilising and reacting to information about their environment (Shettleworth, 2010). Cognition is a versatile trait, enabling wildlife to adapt their behaviours to meet environmental challenges (Cauchoix et al., 2018). In urban environments, where humans are a dominant feature (Gross, 2016), human presence and activity are expected to induce significant selective pressures on various aspects of wildlife (Lerman et al., 2020; Schell et al., 2021). However, which characteristics of human activity affect wildlife cognitive performance remains largely unclear even though such investigations are crucial for sustainable urban planning and city management, particularly in designing green spaces for recreational use (Cardona et al., 2024).

Human activities may provide opportunities for wildlife to consume anthropogenic or novel food via humans directly feeding them or indirectly from feeding stations, waste bins, and food left behind from human gatherings (e.g., (Belant, 1997; Burt et al., 2021; Demeny et al., 2019; L. L. Griffin & Ciuti, 2023; Murray et al., 2015; Rimbach et al., 2023), thus increasing the likelihood of wildlife settling in urban habitats (Goumas et al., 2020; Wilson et al., 2020). However, human activities can often disturb (cause behavioural changes to) urban wildlife (Blanc et al., 2006). Wildlife’s behaviour can be affected by the characteristics of human activities such as the type of activities, the number of humans present in an area, the distance from potential threats, and the speed of the threat stimulus (Bateman & Fleming, 2014; Blanc et al., 2006; Burton et al., 2024; Ramellini et al., 2024; Tablado & Jenni, 2017). For example, urban hooded crows (*Corvus cornix*), Eurasian robins (*Erithacus rubecula*), and wood pigeons (*Columba palumbus*) adjust their activity patterns depending on the number of humans in a park (Ramellini et al., 2024), and Eastern grey squirrels (*Sciurus carolinensis*) show higher alertness in areas with high human activity, particularly where dogs are present (Cooper et al., 2008). Adverse impacts can result from common, seemingly harmless activities, such as walking, dog walking, and cycling. Disturbance may cause both short- and long-term effects on wildlife activities, leading them to adjust their spatial usage and temporal activity patterns (Burt et al., 2021; Doherty et al., 2021; Gaynor et al., 2018), which can have fitness consequences (Bötsch et al., 2017). Despite this, investigations on the impacts of various human activities on wildlife behaviour and, in particular, cognition remain limited.

Investigations into urban wildlife cognition are still in the early stages (Lee & Thornton, 2021) but a few human-dependent factors have already been shown to affect wildlife cognition, in particular innovation. Innovation relies on cognition (Ducatez et al., 2015), as individuals can use existing behaviours to overcome obstacles and obtain food rewards from a novel artificial apparatus (e.g., feeders or puzzle box) (Kummer & Goodall, 1985; Reader & Laland, 2003). Innovative problem-solving ability can be assessed using an artificial food-extraction paradigm (A. S. Griffin & Guez, 2014). Thus far, the few studies that have examined the effects of specific human-related factors on innovation indicate that the presence of a human can directly affect the ability of wild-caught captive juvenile house finches (*Haemorhous mexicanus*) to extract food from a novel artificial feeder (Cook et al., 2017) and that the intensity of humans directly affects population level solving success and individuals’ solving latency in wild Eurasian red squirrels (*Sciurus Vulgaris*) (Chow et al., 2021). Human presence has also been shown to indirectly affect other cognitive abilities, such as generalisation or the capacity to apply learned information to solve similar yet different problems (Chow et al., 2024). However, more focus is needed on comparing the diverse human-related factors. Such research can identify specific features of human activity that shape urban wildlife cognition, which can help to identify the traits and mechanisms that enable urban wildlife to adapt to or thrive in urban environments.

Here, our goal was to examine how different aspects of human activity affect the innovative problem-solving performance of urban Eurasian red squirrels. The ability to innovate is likely related to the fitness of squirrels. Red squirrels demonstrate a remarkable ability to adapt to varying urban environments (Babińska-Werka & Żółw, 2008; Jokimäki et al., 2017; Shimamoto et al., 2020; Thomas et al., 2018; Uchida et al., 2016) and take advantages from living alongside humans (Fingland et al., 2022; Wist et al., 2022). They are opportunistic foragers and food and habitat generalists (Reher et al., 2016; Thomas et al., 2018); they consume anthropogenic food and extract food from novel artificial problems (e.g., (Chow, Clayton, et al., 2021; Chow et al., 2018)).

We used an established food-extraction task (hereafter, puzzle box) to test innovative problem-solving in red squirrels from different populations (Chow et al., 2018; Chow et al., 2021). These previous works showed that increased exposure to humans led to fewer urban red squirrels succeeding (population level), but caused those squirrels that did succeed to have faster solving times over successes (individual level). However, it remains unclear what aspects of human activities influence squirrels’ innovative problem-solving performance. To address this gap, we conducted a field experiment with urban red squirrels, relating their innovative problem-solving performance to three key aspects of human activities: number of humans, type of human activity, and distance to the nearest footpath.

Our general hypothesis was that the three aspects of human activity would affect squirrels’ innovative problem-solving performance, both at the population and individual level, due to creating disturbance. Below, we outline the rationale and predictions for each specific variable:

1. *Number of humans present.* The number of humans present in an area reflects the level of activity intensity in that area. We followed previous research (Chow et al., 2021) and hypothesised that an increased number of humans in a site (i.e., higher activity intensity) would affect the innovative problem-solving performance of squirrels. However, if human activity intensity is consistent in an area, urban wildlife may habituate to human presence as it becomes predictable, thus reducing the need to flee (MacArthur et al., 1979). In such cases, human presence may not affect problem-solving performance.
2. *Types of human recreational and leisure activity.* This variable captures characteristics of human activity at a more granular level. We predicted that any type of human activity would negatively affect squirrels’ innovative problem-solving performance. Urban red squirrels have been shown to initiate flight responses to approaching humans or dogs (Uchida et al., 2019), so we expected walking or dog walking would negatively affect squirrels’ innovative problem-solving performance. We further included cycling and playground activities to compare the varied impacts on solving performance.
3. *Distance to the nearest footpath.* Footpaths are designated areas for human activity. Previous studies have shown that close proximity to these paths can influence wildlife behaviour, for example, increased vigilance in elks (*Cervus elaphus*) and flushing behaviour in vesper sparrows (*Pooecetes gramineus*) and western meadowlarks (*Sturnella neglecta*) (Ciuti et al., 2012; Miller & Knight, 2001). We therefore hypothesised that a shorter distance to a footpath would negatively affect squirrels’ innovative problem-solving performance due to increased exposure to a perceived threat or stressor. Alternatively, the proximity to footpaths may not significantly influence squirrels’ performance because they have become habituated to footpaths as a result of predictable indicators of human activity (see the first hypothesis).

## Materials and Methods

### Study sites

Between August and November 2021, we conducted a field experiment in 15 green spaces (‘sites’) in Oulu, Finland (see map on Google Earth). These sites are fragmented urban forests and urban parks that are defined by the city management of Oulu and via Google Maps. The sites are mostly flat and have footpaths designated by city management. According to the Finnish Public Order Act (Järjestyslaki 27.6.2003/612, Section 14), dogs must be on leash in public areas. Despite this, our observations and records indicate that dogs are frequently unleashed. In all sites, there were trampling impacts on the ground, indicating humans (and their dogs) frequently traversed areas amongst the trees and bushes off the main trails. In all sites, there were no feeders for squirrels and we did not observe humans feeding squirrels. A range of common human activities were observed in these areas, including walking, dog walking, playground activities, and cycling. Activity levels varied across sites (Table 1). The green spaces in Oulu primarily contain forested sections with a mix of deciduous and coniferous trees, including silver birch (*Betula pendula*), downy birch (*B. pubescens*), Norway spruce (*Picea abies*), and Scots pine (*Pinus sylvestris*), creating a blend of natural habitat within the urban environment and providing safety and food sources for red squirrels (Rubino et al., 2012). We avoided sampling the same individuals at different sites (i.e., pseudo-replication) by choosing locations that were 400-500 m apart, which is the average movement of urban squirrels (Andrén & Delin, 1994). Individual identity was confirmed via video recordings (see below) and no squirrels were seen visiting multiple sites throughout the data collection period. In each site, we chose a location that was safe for squirrels for the experiment (e.g., close to a tree, under a tree canopy and away from major roads) to minimise potential fatalities (Fingland et al., 2022). The distance between the puzzle box location at each site and the nearest footpath varied (15-79 m, table 1). We attached a trail camera (Browning model no: BTC-7E-HP4) to a tree to monitor squirrels’ task performance and for individual identification. We followed the protocol by Chow and colleagues (Chow et al., 2018; Chow et al., 2021), identifying each individual using their unique characteristics (e.g., facial features, fur colour, tail shapes) from video footage (see supplementary materials Note S2). This process involved an initial intensive frame-by-frame analysis of video footage to identify each squirrel followed by reanalysing the footage to re-identify the squirrel at different times (Cohen’s Kappa intra-reliability result = 0.99).

**Table 1.**
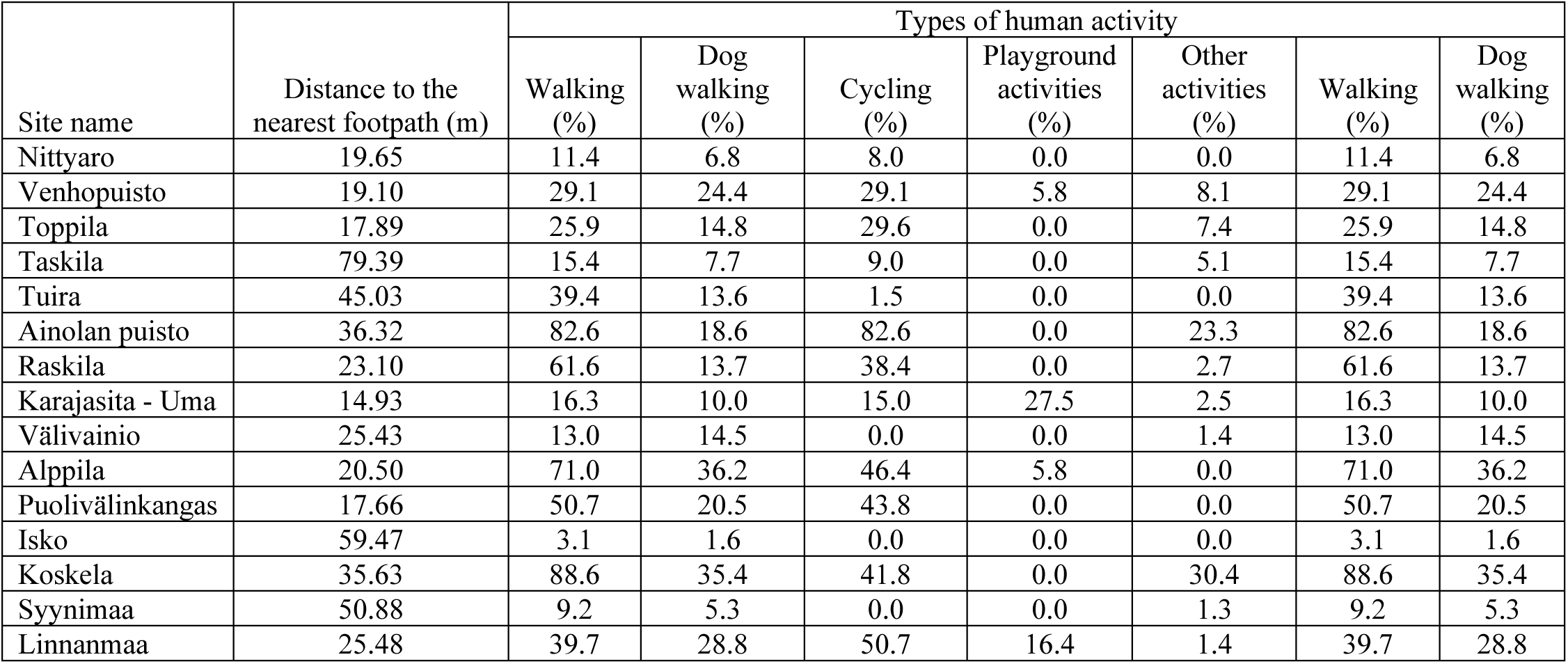
Information about each site (N = 15). The table includes the site name, distance to the nearest footpath in meters (m), and the number of recordings for each type of human activity in percentage. Types of activities include walking, dog walking, cycling, playground activities, and other activities (e.g., bin collection, grass cutting, exercising, and snow-skating). Percentage is calculated from the number of records of each type of activity divided by the total number of records across the field experimental day in a site.

### Puzzle box

The puzzle box presented to squirrels was adapted from an established task that has been solved by ∼ 60% of red squirrels from different populations (Chow et al., 2018; Chow et al., 2021), thus making it a suitable task to reveal variations of performance for this species. We adapted the task by increasing the number of levers (and nuts) in the box to help reduce how often we needed to visit/check each site (i.e. less refilling was required which minimised our disturbance during the field experiment). The puzzle box was made of transparent Plexiglas sheets (shown in Figure 1 and inserts with dimensions). It contained nuts (hazelnut kernels) each positioned within a hole of a lever on top of an immobile plank. Only half of the levers (12 of 24) contained a nut. Squirrels could see and smell the nuts through the box given the various small holes, but could not directly reach the nuts. To solve the task, a squirrel had to either push a lever if the nut was on the squirrel’s side of the box, but had to pull a lever if the nut was on the other side of the box (Figure 1a, video S1). These actions would cause the hole in the lever around the nut to align with a hole in the plank, thus allowing the nut to fall, hit the slanted floor, and roll out from the box from one of the side openings (22.5 cm x 3 cm).

**Figure 1.**
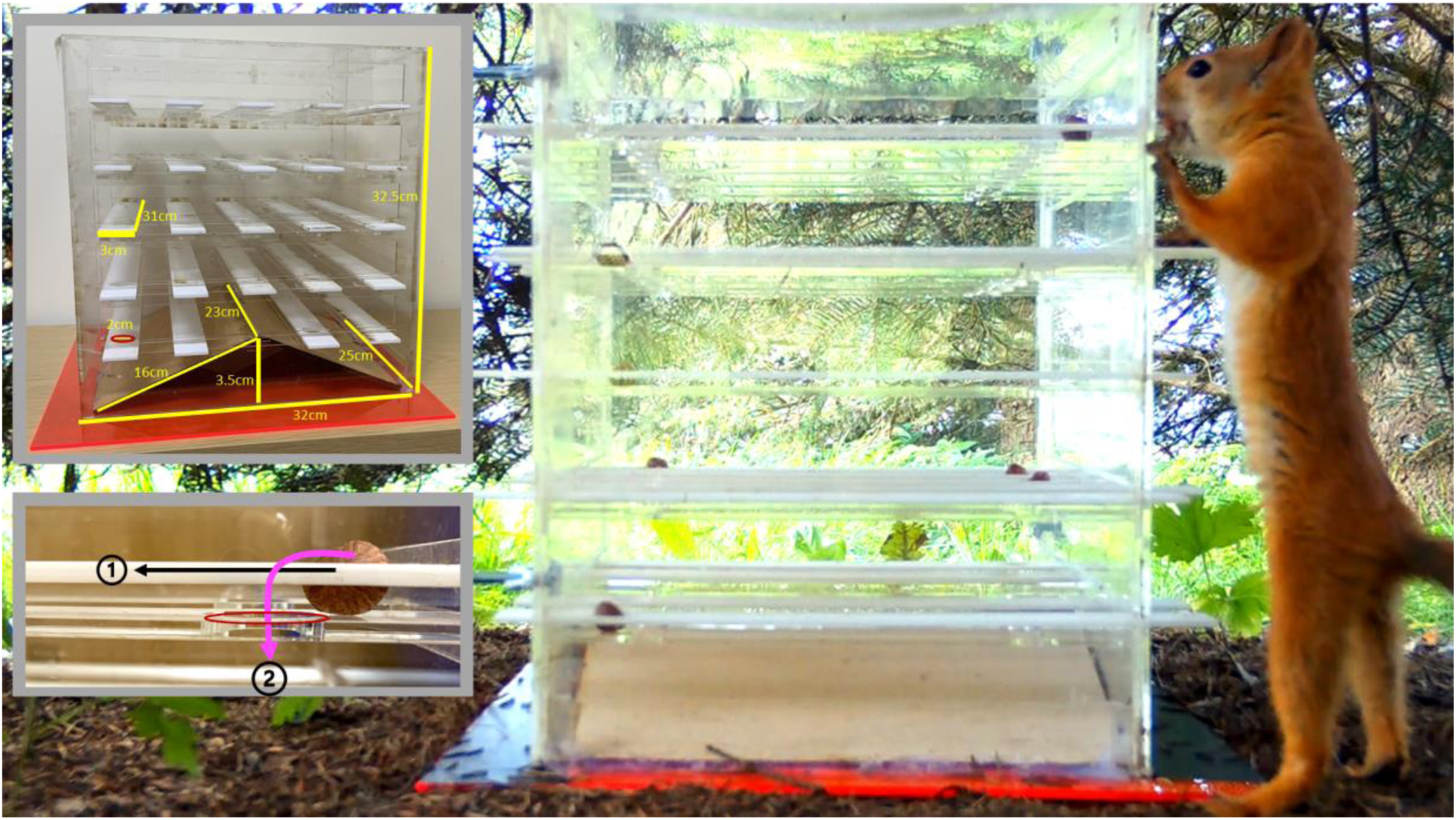
The puzzle box used in the experiments required innovation from squirrels to obtain a highly desired food. The main image shows a squirrel (Bis Bis) attempting to solve the novel food-extraction task used in this study (Dimensions shown in upper left inset). Squirrels were required to push a lever away from themselves if the nut on that lever was positioned near the side where the squirrel was. However, if the nut on a lever was positioned near the opposite side of the squirrel, they had to pull the lever. These actions would cause two holes (one in the lever and one in an immobile plank under the lever) to align and allow the nut to fall through and out of the puzzle box (inset bottom left, 1-2).

### Procedures

The whole experiment in each site lasted for an average of 18 days. Before the puzzle box was presented to squirrels, we baited the location twice a day using unshelled hazelnuts; this aimed to attract squirrels to regularly visit the location. A trail camera was mounted to a tree, around 1 m away from the box to capture squirrels’ visitation, and allowed us to identify individual characteristics and behaviour throughout the experiment. Once squirrels began visiting and obtaining nuts from the location regularly, we carried out a habituation phase. In this phase, we presented the box without levers to squirrels for 4-5 hours and placed hazelnuts around the puzzle box; this phase aimed to minimise any neophobic response to the box. When the squirrels obtained all the nuts around the box, we started the main experiment in which levers were inserted into the box. Data were collected daily in each site from dawn to noon (for an average of 10 consecutive days, ranging from 8 to 12 days), during squirrels’ most active time regardless of weather conditions; this aimed to minimise weather as a confounding variable. The squirrels were free to come and leave the location to ensure their motivation (for food) was high for the experiment. During the experiment, we checked and refilled the box 3-4 times per day, with an inter-check interval of 45 minutes - 1.5 hours; this varied time interval aimed to minimise social interference and allow subordinates to participate in the experiment (less than 2% of the video clips had two or more squirrels). In each check, we randomised the facing direction of the box, the levers that contained a nut, and the facing direction of the levers.

## Measurements

### Human activity factors

We obtained three records to measure human-related factors including:

1. Human presence nearby was recorded using the number of humans within 100 metres (in radius) around the box, which reflects the disturbance threshold distance noted for small mammals (Dertien et al., 2021). Four to five records were made per day, including a record before setting up the box, a record during each check of the box (2-3 per day), and a record at the end of the field experiment for the day. Each record entailed a 5-minute observation of the immediate area around the site. For each site, we summed the number of humans across all records and divided this total by the number of records made across the field experiment days at that site to obtain the mean number of humans present per record.
2. Types of human activity, recorded using *ad-hoc* recording during the 5-minute observation. Five categories included ‘walking’, ‘dog walking’, ‘cycling’, ‘playground activities’ (e.g., gathering, playing in the playground, ball games such as football), and ‘other activities ’ (e.g., construction work, bin collection, grass cutting). During each observation, we noted the number of humans engaged in that activity. For each type of human activity, the total number of individuals recorded was used to determine the frequency of that activity. To calculate the mean frequency for each activity per record, we divided the total number of humans participating in each activity by the total number of records across the field experiment days at each site. This approach allowed us to estimate the intensity of human presence with a more specific focus on the type of activity, in contrast to the general human presence calculated previously.
3. Distance to the nearest footpath was measured as the shortest distance (in metres) between the nearest footpath and the location where we placed the puzzle box. To obtain this record, we dropped a pin on the ground at the box location and used a measuring tape to calculate the shortest distance to the footpath. We further confirmed the distance again on Google Maps; the location of the box was pinned on Google Maps, followed by measuring the distance between the location and the nearest footpath three times, and we took the average distance (m) for this measure.

### Innovative problem-solving performance

The innovative problem-solving performance of the squirrels was assessed through their interaction with a lever in the food-extraction task. Following (Chow et al., 2021), when a squirrel manipulated a lever using its paws, teeth or nose, we obtained two records until the squirrel ceased its attempts. The first record was ‘solving outcome’, which indicated whether the manipulation was successful, i.e.caused a nut to fall or an empty lever to be dislodged (these two scenarios were considered successful to account for various motivations, e.g. hunger or play). The second record noted the ‘solving latency’, which was the time spent manipulating the lever. In line with previous studies (Chow et al., 2021), squirrels were classified based on the solving outcome: those who solved the problem multiple times were categorised as ‘innovators’ and those who only succeeded once but could not replicate the success or failed to solve the problem throughout the presentation period were ‘non-innovators’. Among the innovators, we further considered squirrels that solved the problem on their first visit as ‘first-visit solvers’ and those that solved the problem on subsequent visits as ‘subsequent solvers’. To obtain population-level performance, we counted the number of squirrels that successfully solved the problem in each site and divided this number by the total number of squirrels participating in the problem (i.e., the first-visit success rate at the population/site level). Only squirrels that had successfully solved the problem provided data for solving latency when analysing individual-level performance. For each solver, we summed across all the durations of failed manipulations until a success occurred to obtain the first-success latency.

#### Data analyses

All data were analysed using R version 4.4.1 (*The Comprehensive R Archive Network*, n.d.). Before running the models, we first checked the correlations among variables of interest using Pearson’s *r* (see table S1-S4). To avoid multicollinearity that would induce interpretation problems (Allen, 1997), we included variables that did not have moderate to high correlation *r* ≥ 0.5 (Dancey & Reidy, 2007). After each model was run, we used the Variance Inflation Factor (VIF) in the ‘car’ package (Fox et al., 2012) to further assess multicollinearity (VIF < 5, tolerance > 0.2) (see results below). All results reported here are two-tailed and the significance level was set at p ≤ 0.05.

The proportion of success at the population (site) level was analysed using beta regression of the ‘betareg’ package (Cribari-Neto & Zeileis, 2010). The first-success latency at the individual level was analysed using a Generalized Linear Mixed Model (GLMM) with gamma log-link distribution in the ‘glmmTMB’ package (Magnusson et al., 2017), which can accommodate the skewed distribution. In all GLMMs, random variables were site and squirrel identity where possible.

We first examined the factors that affected the innovative problem-solving performance of first-visit solvers (N= 43). The two main models included the fixed effects of distance (m) to the nearest footpath and the mean number of humans nearby whereas the proportion of success at the population level was the response variable in one model, and the first-success latency at the individual level in the other model. In both models, we added the squirrel population (the number of squirrels that participated in the experiment at each site) as another fixed effect because this variable has been shown to affect solving performance (Chow et al., 2021). We then reran the models by including the subsequent solvers (N = 10) to examine the factors that affected the innovative problem-solving performance of all innovators (N = 53).

To examine whether the types of human activity and distance to the nearest footpath affected each problem-solving performance measure, we ran four models due to the high correlation between walking and cycling (*r* = 0.94 & *r* = 0.95) (see Table S2 & 4). Each problem-solving performance measure had two models. One model included fixed factors of walking, dog walking, playground activities and distance of nearest footpath. The other model included the following fixed factors: cycling, dog walking, playground activities and distance of nearest footpath. In all the models, variables were standardised to compare their effect size.

## Results

### Human presence nearby, distance to the nearest footpath, squirrel population and solving success

In the selected 15 urban green spaces, a total of 64 squirrels participated in the food-extraction problem. 53 (83%) squirrels were innovators, with 43 (68%) squirrels solved the problem on their first visit (i.e., first-visit solvers) and another 10 squirrels solved the problem on a subsequent visit (i.e., subsequent solvers). At the population (site) level, human presence nearby, distance to the nearest footpath and squirrel population explained 18% of the variance in the proportion of first-visits success (Table 2 Model 1a, Figure 2a-c). First-visit success rate significantly decreased with an increased mean human presence nearby (Z = -1.98, P = 0.048) (Figure 2a). However, neither squirrel population (Z = -0.13, P = 0.894) (Figure 2b) nor distance to the nearest footpath (Z = -0.37, P = 0.710) (Figure 2c) affected the first-visit success rate. Another model including all innovators (first-visit or subsequent-visit solvers) showed that human presence nearby, distance to the nearest footpath and squirrel population explained 29% of the variance in the proportion of success at the population level (Table 2 Model 1b). Similar to the model of the first-visit success, mean human presence nearby was the only factor that significantly negatively affected the overall success rate at the population level (Z = -2.60, P = 0.009).

**Figure 2.**
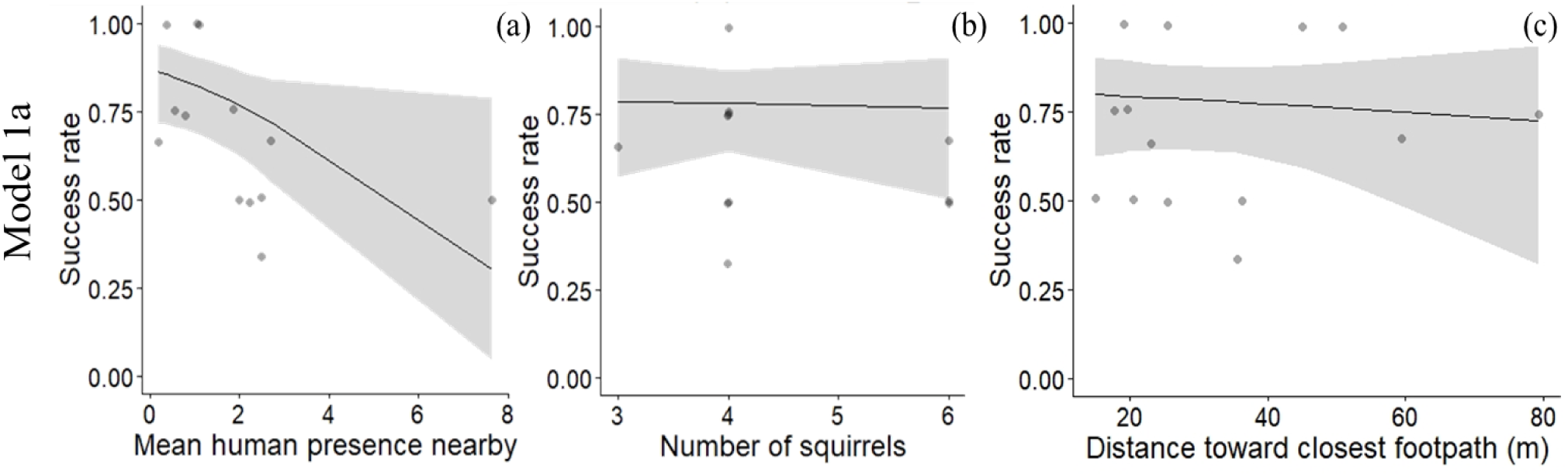
Figure of Model 1a, showing the effects of (a) mean human presence nearby, (b) squirrel population, and (c) distance to the nearest footpath (m) on the first-visit success at the population/site level (N = 15). Human presence nearby was the mean number of humans present within a 100 m radius around the puzzle box during each check. Squirrel population was the number of squirrels that participated in the experiment. Distance to the nearest footpath (m) was the shortest distance between the puzzle box and the nearest footpath.

**Table 2.** Models (M) included fixed factors of human activities to predict success rate at the population level of (a) the first-visit solvers (N = 43) and (b) first and subsequent solvers (N = 53). Model 1 examined general factors that included squirrel population, mean number of human presence nearby and distance to the nearest footpath as fixed factors. Models 2 and 3 emphasised the effects of the types of human activities on innovative problem-solving success at the population/site level. Model 2 included walking, dog walking, playground activities and distance to the nearest footpath as fixed factors. Model 3 included cycling, dog walking, playground activities and distance to the nearest footpath as fixed factors. Squirrel population was the number of squirrels that participated in the study at each site. Human presence nearby was the mean number of humans within 100 metres around the puzzle box during each check. The nearest distance footpath was the actual distance (cm) from the box to the edge of the nearest footpath. Human activities included walking, dog walking, cycling, and playground activity. This table shows standardised estimate (Est), standard error (S. E.), Z and P values, and Marginal R(MR^2^). Bold values indicate *P* < 0.05.

| M | Fixed factors | Response variables |  |  |  |  |  |  |  |  |  |
| --- | --- | --- | --- | --- | --- | --- | --- | --- | --- | --- | --- |
|  |  | (a) Proportion of first-visit success<br>(first-visit solvers) (N = 43) |  |  |  |  | (b) Proportion of first and subsequent<br>visit success (all solvers) (N = 53) |  |  |  |  |
|  |  | Est | S.E. | Z | P | MR<sup>2</sup> | Est | S.E. | Z | P | MR<sup>2</sup> |
| 1 | Squirrel population | -0.04 | 0.32 | -0.13 | 0.894 | 0.18 | 0.06 | 0.27 | 0.20 | 0.838 | 0.29 |
|  | Mean number of human presence per record | -0.66 | 0.33 | -1.98 | **0.048** |  | -0.43 | 0.17 | -2.60 | **0.009** |  |
|  | Distance of nearest footpath | -0.12 | 0.32 | -0.37 | 0.710 |  | -0.01 | 0.02 | -0.87 | 0.385 |  |
| 2 | Walking | -0.58 | 0.29 | -2.02 | **0.043** | 0.29 | -0.71 | 0.25 | -2.79 | **0.005** | 0.49 |
|  | Dog walking | -0.71 | 0.31 | -2.25 | **0.024** |  | -0.37 | 0.29 | -1.28 | 0.200 |  |
|  | Playground activities | -0.79 | 0.29 | -2.71 | **0.007** |  | -0.80 | 0.26 | -3.01 | **0.002** |  |
|  | Distance of nearest footpath | -0.58 | 0.32 | -1.83 | 0.067 |  | -0.49 | 0.30 | -1.65 | 0.099 |  |
| 3 | Cycling | -0.46 | 0.28 | -1.67 | 0.095 | 0.25 | -0.59 | 0.25 | -2.38 | **0.017** | 0.44 |
|  | Dog walking | -0.76 | 0.32 | -2.40 | **0.017** |  | -0.42 | 0.29 | -1.45 | 0.148 |  |
|  | Playground activities | -0.72 | 0.29 | -2.49 | **0.013** |  | -0.72 | 0.26 | -2.73 | **0.006** |  |
|  | Distance of nearest footpath | -0.66 | 0.32 | -2.05 | **0.041** |  | -0.59 | 0.30 | -1.95 | 0.051 |  |

### Types of human activity, distance to the nearest footpath, and solving success

The most common human activity across the urban green spaces was walking (37.1%), followed by cycling (26.4%), dog walking (16.8%), other activities (5.6%) and playground activities (3.7%) (see Table 1 for human activity by site). The model that included walking, dog walking, playground activities and distance to the nearest footpath explained 29% of the variance in first-visit success at the population level (N = 43, Table 2 Model 2a). Except for the distance to the nearest footpath, all types of human activity significantly decreased the first-visit success rate at the population level (Table 2 Model 2a, Figure 3 Model 2a). Playground activities had the highest effect size (-0.79) followed by dog walking (-0.71) and walking (-0.58). When including first and subsequent solvers in the model (N = 53), the factors explained 49% of the variance in the success rate at the population level. Walking and playground activities significantly affected the success rate at the population level (Table 2 Model 2b). Walking had the highest effect size (-0.71) followed by playground activities (-0.80).

**Figure 3.**
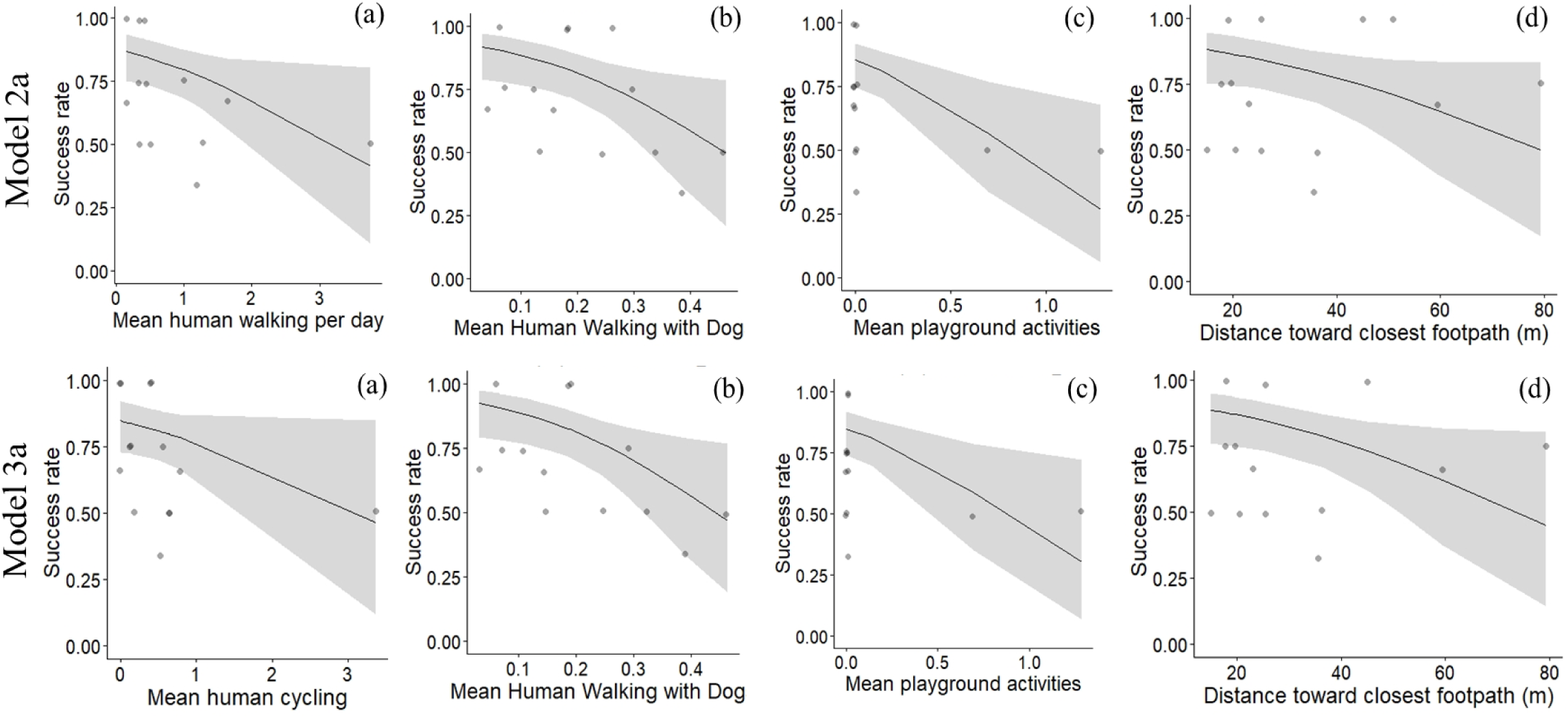
Effects of different types of human activities (a-c) and distance to the nearest footpath (d) on the first-visit success rate at the population/site level (N = 15). Model 2a includes (a) walking, (b) dog walking, (c) playground activities and (d) distance to the nearest footpath. Model 3a includes (a) cycling, (b) dog walking, (c) playground activities and (d) distance to the nearest footpath.

The model that included cycling, dog walking, playground activities and distance to the nearest footpath for first-visit solvers explained 25% of the variance in first-visit success at the population level (Table 2 Model 3a). Except for cycling, all the variables significantly decreased the likelihood of first-visit success (Table 2 Model 3a, Figure 3 Model 3a). Dog walking (-0.76) had a slightly higher effect size than playground activities (-0.72). When including first and subsequent solvers (all innovators) in the model, the factors explained 44% of the variance in success rate at the population level. Cycling and playground activities significantly affected the success rate at the population level (Table 2 Model 3b). Playground activities had the highest effect size (-0.72) followed by cycling (-0.59).

### Human presence nearby, distance to nearest footpath and first-success latency

First-visit solvers (N = 43) took around 6 seconds (SD = 7.01) to cause a nut fall or to dislodge an empty lever (see Panda, an innovator in the <u>video</u> S1). For these first-visit solvers, the model that included squirrel population, mean human presence nearby, and distance to the nearest footpath (m) explained 17% of the variance in first-success latency (Table 3 Model 1a, Figure 4a-c). Only human presence nearby significantly affected these innovators’ first-success latency (Z = -2.20, P = 0.028); increased mean human presence nearby decreased their first-success latency (Figure 4a). Ten more squirrels subsequently solved the problem (N = 53), with a mean solving latency of 8 seconds (SD = 9.50). Squirrel population, mean human presence nearby, and the distance to the nearest footpath explained 17% of the variance in their first-success latency (Table 3 Model 1b). Shorter first-success latency was associated with a larger squirrel population (Z = 1.98, P = 0.048) and a higher mean human presence nearby (Z = -2.20, P = =0.028). Distance to the nearest footpath did not affect the first-success latency (Z = -0.11, P = 0.916).

**Table 3.** Models (M) included fixed factors of human activities to individual first-success latency of (a) first-visit solvers (N = 43) and (b) all solvers (N = 53). Model 1 examined general factors that included squirrel population, mean number of humans present nearby and distance to the nearest footpath as fixed factors. Models 2 and 3 emphasised the effects of the types of human activities on innovative problem-solving success at the population/site level.

|  |  | Response variables |  |  |  |  |  |  |  |  |  |
| --- | --- | --- | --- | --- | --- | --- | --- | --- | --- | --- | --- |
|  |  | (a) first success latency for first-visit solvers (N = 43) |  |  |  |  | (b) first-success latency for all solvers (N = 53) |  |  |  |  |
| M | Fixed factors | Est | S.E. | Z | P | MR<sup>2</sup> | Est | S.E. | Z | P | MR<sup>2</sup> |
| 1 | Squirrel population | 0.18 | 0.20 | 0.92 | 0.358 | 0.17 | 0.31 | 0.16 | 1.98 | **0.048** | 0.17 |
|  | Mean number of human presence per record | -0.39 | 0.18 | -2.20 | **0.028** |  | -0.42 | 0.19 | -2.20 | **0.028** |  |
|  | Distance of nearest footpath | 0.11 | 0.15 | 0.72 | 0.472 |  | -0.02 | 0.15 | -0.11 | 0.916 |  |
| 2 | Walking | -0.03 | 0.19 | -0.14 | 0.887 | 0.30 | -0.22 | 0.19 | -1.14 | 0.256 | 0.20 |
|  | Dog walking | -0.66 | 0.22 | -2.96 | **0.003** |  | -0.21 | 0.16 | -1.29 | 0.196 |  |
|  | Playground activities | -0.29 | 0.20 | -1.44 | 0.151 |  | 0.20 | 0.16 | 1.26 | 0.208 |  |
|  | Distance of nearest footpath | -0.14 | 0.18 | -0.77 | 0.441 |  | 0.14 | 0.16 | 0.85 | 0.397 |  |
| 3 | Cycling | 0.01 | 0.19 | 0.06 | 0.950 | 0.30 | -0.19 | 0.20 | -0.96 | 0.338 | 0.20 |
|  | Dog walking | -0.68 | 0.22 | -3.07 | **0.002** |  | -0.23 | 0.16 | -1.44 | 0.151 |  |
|  | Playground activities | -0.28 | 0.20 | -1.43 | 0.152 |  | 0.24 | 0.16 | 1.51 | 0.130 |  |
|  | Distance of nearest footpath | -0.14 | 0.18 | -0.79 | 0.427 |  | 0.14 | 0.16 | 0.83 | 0.404 |  |

**Figure 4.**
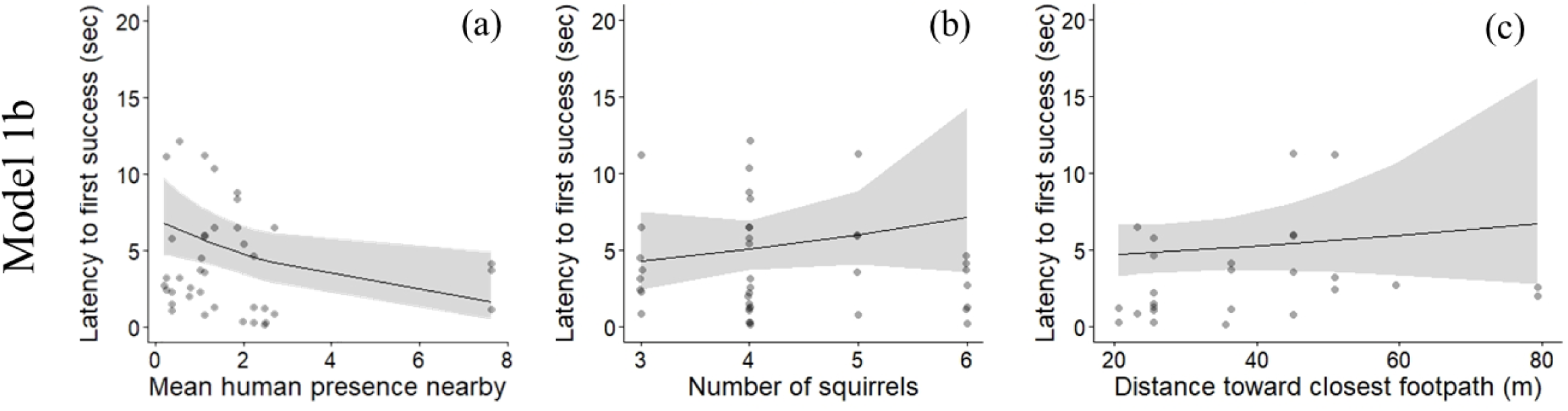
Figure of Model 1b, showing effects of (a) mean human presence nearby, (b) squirrel population, and (c) distance to the nearest footpath (m) on the first-success latency of first-visit solvers (N = 43). The human presence nearby included the mean number of humans within a 100 m radius around the puzzle box during each check. Squirrel population was the number of squirrels that participated in the problem-solving task at each site. Distance to the nearest footpath was the shortest distance between the location of the puzzle box and the footpath (m).

### Types of human activity, distance to the nearest footpath, and solving latency

For the first-time solvers (N = 43), the model that included walking, dog walking, playground activities, and the distance to the nearest footpath explained 30% of the variance in the first-success latency (Table 3 Model 2a). Only dog walking significantly affected their first-success latency (Z = -2.96, P = 0.003) (Table 3 Model 2a, Figure 5 Model 2b). The model that included cycling, dog walking, playground activities and distance to the nearest footpath also explained 30% of the variance in the first-success latency (Table 3 Model 3b). Similar to the previous model, only dog walking affected first-success latency (Z = -3.07, P = 0.002). In both models, more dog walking led to shorter first-success latency (Figure 5 Model 3b). For both first and subsequent solvers (N = 53), both models explained 20% of the variance in first-success latency and none of the activities nor distance to the nearest footpath significantly affected first-success latency (Table 3 Model 2b & Model 3b).

**Figure 5.**
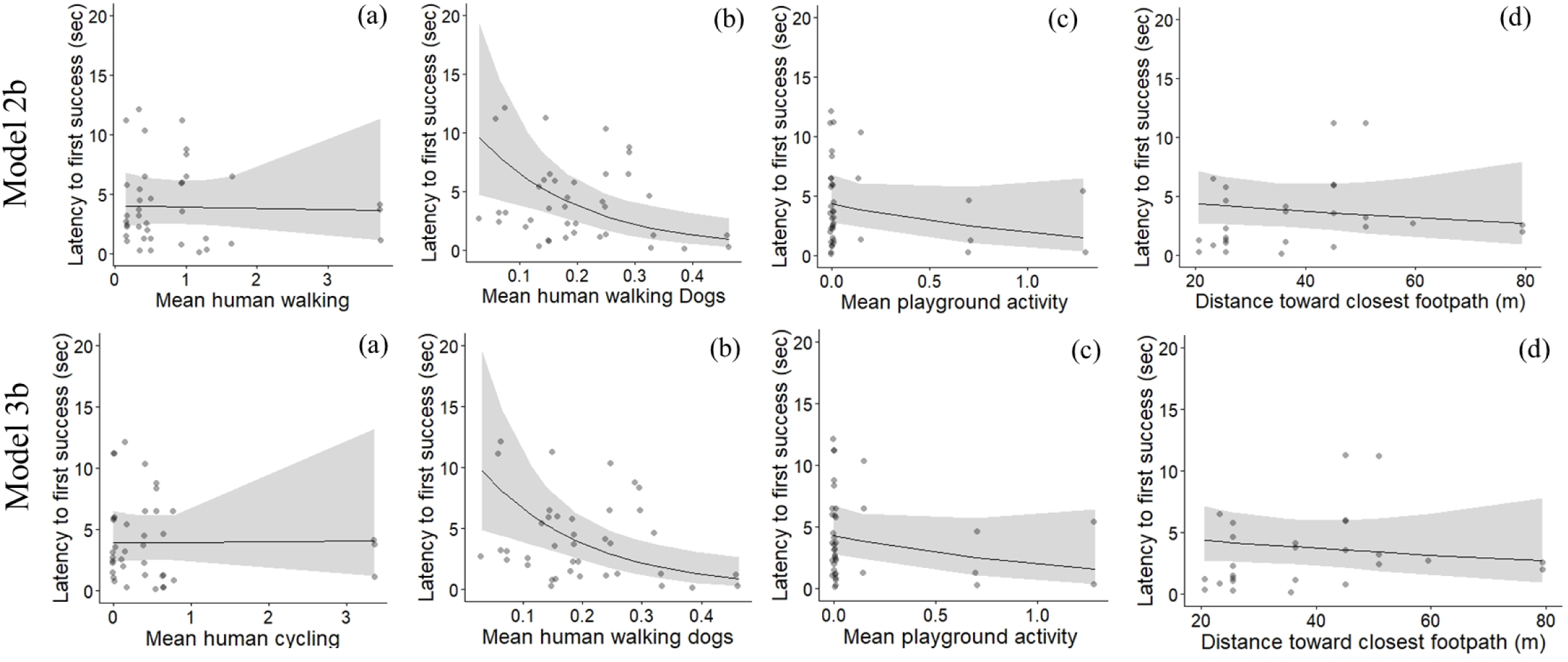
Models 2 and 3 examined the effects of different types of human activities (a-c) and distance to the nearest footpath (d) on first-visit solvers’ first-success latency (N = 43). Model 2b included (a) walking, (b) dog walking, (c) playground activities, and (d) distance to the nearest footpath. Model 3b included (a) cycling, (b)dog walking, (c) playground activities, and (d) distance to the nearest footpath.

Model 2 included walking, dog walking, playground activities and distance to the nearest footpath as fixed factors. Model 3 included cycling, dog walking, playground activities and distance to the nearest footpath as fixed factors. Squirrel population was the number of squirrels that participated in the study at each site. Human presence nearby was the mean number of humans within 100 metres around the puzzle box during each check. The nearest distance footpath was the actual distance (cm) from the puzzle box to the edge of the nearest footpath. Human activities included walking, dog walking, cycling, and playground activity. This table shows standardised estimate (Est), standard error (S. E.), Z and P values, and Marginal R(MR^2^). Bold values indicate *P* < 0.05.

### Solving performance and interaction between types of human activities and distance to the nearest footpath

For each performance measurement, two additional models were run to examine only the interaction effects of the types of human activity and distance to the nearest footpath. The two models of first-visit success at the population explained 8-10% of the variance and showed no significant interaction effect on first-visit success at the population level (Table 4 Model 1-2a). After including subsequent solvers in the model, the model that included interactions of walking and distance to the nearest footpath remained the same (Table 4 Model 1b); there was no significant interaction effect detected in the model. For the model that included the interaction of cycling and distance to the nearest footpath, the interaction effects between all types of activities and distance to the nearest footpath were significant (Table 4 Model 2b). Both models of first-success latency of first-time solvers explained 10% of the variance and had no significant interaction effects on first-success latency (Table 4 Model 3-4a). Similar results were obtained when all solvers were included in the model and the model explained 14-17% of the variance of first-success latency (Table 4 Model 3-4b).

**Table 4.** Interaction effects of types of human activity and distance to the nearest footpath for (a) first-visit success at the population level (N=15) and (b) first-success latency of innovators at the individual level (N=43). These models only included interaction effects. This table shows standardised estimate (Est), standard error (S. E.), Z and P values, and Marginal R(MR^2^).

|  |  | Response variables |  |  |  |  |  |  |  |  |  |
| --- | --- | --- | --- | --- | --- | --- | --- | --- | --- | --- | --- |
|  |  | Success at the population level (N =15) |  |  |  |  |  |  |  |  |  |
|  |  | (a) first-visit success rate |  |  |  |  | (b) first and subsequent visit success rate |  |  |  |  |
| M | Fixed factors | Est | S.E. | Z | P | MR<sup>2</sup> | Est | S.E. | Z | P | MR<sup>2</sup> |
| 1 | Walking*footpath | 0.97 | 1.00 | 0.97 | 0.332 | 0.10 | 0.30 | 0.94 | 0.32 | 0.752 | 0.17 |
|  | Dog walking*footpath | -0.63 | 0.67 | -0.95 | 0.342 |  | 0.06 | 0.62 | 0.09 | 0.927 |  |
|  | Playground activities *footpath | 0.43 | 0.36 | 1.19 | 0.233 |  | 0.64 | 0.33 | 1.93 | 0.053 |  |
| 2 | Cycling*footpath | -0.14 | 1.06 | -1.08 | 0.282 | 0.08 | -1.93 | 0.96 | -2.01 | **0.044** | 0.24 |
|  | Dog walking*footpath | 0.62 | 0.63 | 0.99 | 0.320 |  | 1.27 | 0.57 | 2.21 | **0.027** |  |
|  | Playground activities*footpath | 0.50 | 0.36 | 1.39 | 0.165 |  | 0.75 | 0.31 | 2.43 | **0.015** |  |
|  |  | First-success latency at the individual level |  |  |  |  |  |  |  |  |  |
| M | Fixed factors | (c) first-time solvers (N = 43) |  |  |  |  | (d) first and subsequent solvers (N = 53) |  |  |  |  |
| 3 | Walking*footpath | -0.96 | 0.73 | -1.31 | 0.192 | 0.10 | -0.90 | 0.62 | -1.46 | 0.145 | 0.17 |
|  | Dog walking*footpath | 0.38 | 0.47 | 0.80 | 0.422 |  | 0.35 | 0.36 | 0.96 | 0.337 |  |
|  | Playground activities *footpath | 0.17 | 0.23 | 0.75 | 0.455 |  | -0.28 | 0.18 | -1.54 | 0.125 |  |
| 4 | Cycling*footpath | -0.98 | 0.64 | -1.53 | 0.127 | 0.10 | <-0.01 | 0.01 | -0.37 | 0.171 | 0.14 |
|  | Dog walking*footpath | 0.34 | 0.40 | 0.85 | 0.398 |  | -0.11 | 0.16 | -0.72 | 0.476 |  |
|  | Playground activities*footpath | 0.22 | 0.23 | 0.97 | 0.331 |  | -0.25 | 0.19 | -1.28 | 0.202 |  |

## Discussion

Humans are a main feature of urban habitats and potentially act as a major selective pressure on phenotypic traits including urban wildlife cognition ((Lerman et al., 2020; Schell et al., 2021)). It is crucial to understand the traits that facilitate urban wildlife to adapt to urban environments, which has implications for city planning and management of green space. Here, we conducted a field experiment to examine the impacts of human activities (i.e., human presence, types of activities in urban green space, and distance to the nearest footpath) on urban squirrels’ innovative problem-solving performance. Key findings include: 1) an increased mean human presence nearby negatively affected the success rate at the population (site) level and the first-success latency at the individual level; 2) playground activities and dog walking had a greater impact on first-visit success than other activities; and 3) first-visit solvers were quicker to succeed if dog walking occurred nearby. These results highlight key aspects of human activity affecting squirrels’ innovative problem-solving ability, a trait that is potentially important for successful settlement in urban environments (Barrett et al., 2019).

Previous studies have shown that an approaching human triggers a flight response in squirrels (Uchida et al., 2019, 2020), and increased human intensity around a site has been shown to decrease the success rate of solving a food-extraction task at the population level (Chow et al., 2021). In this study, human presence nearby reflects human intensity at a closer distance (within a 100 m radius of the box). As predicted, the number of humans present nearby significantly affected squirrels’ innovative problem-solving performance. Specifically, increased human presence nearby decreased the first-visit success at the population level, suggesting a negative impact of human presence on squirrels’ innovative problem-solving ability. Nearby humans may disturb squirrels’ attempts to solve problems, causing some squirrels to fail or abandon the task and forage elsewhere after the first attempt. Squirrels that failed their first attempt may try again and succeed in subsequent visits. With increased human presence nearby, the fact that both the first and subsequent solvers showed shorter first-success latency confirmed that human presence and activity are stressors for urban squirrels and that squirrels perceive humans as potential threats (Uchida et al., 2016). During innovation, like that experienced in the current study, squirrels often change from their typical foraging pattern in trees to the ground (Shuttleworth, 2000), which increases their exposure to humans and in turn may intensify squirrels’ perceived stress. Their enhanced solving efficiency in response to increased human presence has been suggested as an adaptive strategy, allowing them to quickly solve the problem and retreat to a tree for safety (Chow et al., 2024).

We sought to determine which specific types of human activities are most impactful. In urban areas, four common types of human activities (walking, dog walking, playground activities, and cycling) likely influence squirrels’ behaviour in response to human disturbance (Bateman & Fleming, 2014; Uchida et al., 2019, 2020). We predicted that human activities like walking and dog walking would negatively affect squirrels’ innovative problem-solving performance. Our results support the prediction, showing that walking and dog walking decreased first-visit success rate at the population level and dog walking decreased first-success latency. One explanation for why walking and dog walking decreased success rate is related to the predictability of these activities (Tablado & Jenni, 2017). Compared with cycling which mostly occurs on the designated footpath, humans (and their dogs) can move freely in these urban green spaces. Their movement can therefore be unpredictable (e.g., going off a footpath, chasing or approaching squirrels) (Weston & Stankowich, 2014), especially since most of our records noted that dogs were not on leash. Despite this, walking has a lower negative impact than dog walking on first-visit success. This result suggests that the squirrels, to some extent, were habituated to humans or may have increased their tolerance in response to human presence or disturbance, which has been shown in another population of urban red squirrels (Uchida et al., 2019, 2020). Nevertheless, urban squirrels have been shown to remain vigilant in response to human walking and walking with dogs (Cooper et al., 2008; Uchida et al., 2019). Remaining vigilant is part of the risk assessment of surrounding environments that can inform individuals when to initiate a flight response (Blumstein, 2003). In our experiment, squirrels may have abandoned problem-solving tasks after detecting humans or dogs nearby. However, their flight response incurs a trade-off with squirrels’ ongoing (foraging) activity, which has been shown in birds in response to humans and with their dogs (Miller & Knight, 2001) and other fitness-related activities (Bötsch et al., 2017). Dog walking, especially when the dog is unleashed, may lead to various adverse effects on wildlife, including injury or death (Weston & Stankowich, 2014). Given that urban squirrels have a high overlapping active time with domestic dogs (Tobajas et al., 2023) and dogs are common predators of urban squirrels (Makowska & Kramer, 2007; Wauters et al., 1997), squirrels may retreat from solving novel problems in response to a perceived increased threat (e.g., human and dogs approaching, see supplementary <u>video</u> S2). The faster solving latency for first-visit solvers suggests one of the strategies that balance foraging and antipredator behaviour is that innovators retreat to a tree and monitor their environment once they have solved the task.

In addition to walking and dog walking, playground activity also negatively impacted first-visit success. Notably, playground activity had the fewest recorded instances among the five types of human activities (see Table 1), but its negative impact on squirrels’ innovation performance was similar to dog walking, a well-established disturbance/threat to wildlife (Weston & Stankowich, 2014) as discussed above. The significance of playground activity to solving success may be related to the characteristics of this type of activity. For example, playground activity, to some extent, is predictable in spatial usage (i.e., in a designated area) but often induces loud noises of varied duration (Rosenthal et al., 2022). Anthropogenic noise may mask the detection of predators or reduce effectiveness in communication, which leads to acute impacts on wildlife behaviour (Blickley & Patricelli, 2010). Because tree squirrels such as the current study species use interspecific and intraspecific calls for threat detection (Randler, 2006), an increase in playground activity likely disturbs their behaviour, especially when they are on the ground attempting to solve problems.

Human activity may provide different cues that wildlife can use as an indicator of threat which can help predict and initiate appropriate responses. For example, herring gulls use the gaze direction of humans to approach food (Goumas et al., 2019) and urban red squirrels use the direction of an approaching human to initiate flight response direction (Uchida et al., 2019). Designated footpaths for humans may also provide high predictability for wildlife. However, our results only partially support the prediction that distance to the nearest footpath independently affects innovative problem-solving. The result that increased distance to the nearest footpath decreased squirrels’ success rate also contradicted our expectations. One possible explanation is related to the intertwined characteristics in urban environments (Boeing, 2018), i.e. the effect of footpaths may depend on other factors like the number of humans, the type of human activity, etc. Our results of the interaction effects show that distance to the nearest footpath and some types of human activity affected the success rate at the population level. Another possible explanation is related to where human activities occur. In urban environments where the activity period of wildlife overlaps with humans, many animals including this species of red squirrels adjust their active time to avoid human peak activity (Stover, 2014). Being further away from a footpath may minimise exposure to humans and dogs, but being nearer to the footpath can increase visibility for early detection of these human-related potential threats. According to the Finnish Public Order Act (Järjestyslaki 27.6.2003/612, Section 14), dogs must be kept on a leash in public areas such as parks and streets. However, our observations indicate more than a third of dog walkers disregarded this law (average 70.6% across all sites). Because many dogs (note that our cameras also caught several domestic cats near the puzzle box) are not on a leash and that humans and dogs can walk freely through the woods, the squirrels may perceive a higher threat further from the footpath. The significant interaction result suggests that some squirrels are able to differentiate different human activities in their risk assessment. Being able to do this with different types of cues has been shown in other tree squirrels species such as fox squirrels, *S. niger*, residing in urban areas differentiating the sound of humans and natural predators (Kittendorf & Dantzer, 2021).

Overall, we demonstrated the impact of human disturbance on an important cognitive trait, innovative problem-solving, in urban red squirrels. Our results highlight the types of human activities, particularly dog walking, that can induce negative effects on squirrels’ ability to solve novel problems. Our findings concerning the negative effects of dogs on squirrel cognition (and overall frequency of dogs off leash) may be useful for urban management and policy decisions, specifically in considering the importance of minimising, and ways to minimise, interactions between pet dogs (and cats) with wild squirrels (and other wild animals). It is expected that these and other human-related activities put a strong selective pressure on wildlife behaviour and cognition, and thereby affect their ability to adapt to urban environments (Sih, 2013). The general results here appear to indicate that urban red squirrel’s innovative problem-solving ability is affected by different factors of human activity. Future studies examining the specific cues that squirrels use in decision-making and risk assessment, and how these cues are related to problem-solving performance, will provide a better understanding of how wildlife adapt to urban environments.

## Acknowledgement

The authors thank the City of Oulu for granting access to the study areas, and A. Chow for logistic support. Biodiversity Anthropocenes for the support, and Fab Lab Oulu for guidance and support in constructing the puzzle boxes.

## Ethics

This experiment did not involve any invasive methods. All individuals were identified using video recordings. The individuals were free to come and leave the task at their own volition. No punishments were administered if they did not participate in the task, or if they failed to solve the puzzle box. The whole procedure adheres to ASAB and ABS guidelines, as well as Finnish law.

## Data availability

All data is uploaded to OSF (DOI 10.17605/OSF.IO/NFTBD) or https://osf.io/nftbd/ (if DOI failed)

