## Supplementary materials for "The Effect of Human Presence and Activity Type on Innovative Problem-Solving of Urban Eurasian Red Squirrels"

[1]Ecology and Genetics Research Units
University of Oulu
Linnamma
Finland 90570

[2]School of Psychology
University of Chester
Chester
UK CH1 4BJ

This document contains additional notes, tables, figures, and links to videos of the main manuscript.
Note S1-S2
Table S1-S4
Video S1-S2

Note S1 Geographic information of the 15 field sites (Oulu, Finland) on Google Earth
https://earth.google.com/earth/d/1vHisjzF8p4AtRtrLEIfn-ANh4tymtODG?usp=sharing

Note S2. Individual identification

To identify each individual and also record the number of squirrels in each site, we used an established method by Chow et al., (2018) using frame-by-frame analysis using Adobe Premiere Pro CS6. The first identification required intensive observer training that lasted for two months. This analysis is similar to the ‘mark-recapture’ and ‘mark-resight’ methods but on video footage. When we saw a squirrel appear on a video for the first time, it was ‘marked’ using their characteristics. Each squirrel was assigned a name and an identification number. This process required back and forth watching different footage so that the individuals’ full characteristics could be revealed from different angles. It was ‘recaptured’ when it reappeared in the subsequent videos. We recorded detailed characteristics of each squirrel that included their facial marking (e.g. a white dot/patch on face), colouration (e.g. orange, burgundy, brown patch on forehead), the colour of their limbs (e.g. orange/dark brown paws or dots on a toe) alongside height (relative to the apparatus), tail and body shape (e.g. full fur tail, half tail). We reidentified the squirrels three to five months after the first identification during which the same coder (PKYC) re-conducted the frame-by-frame analyses of the unmarked individuals. To examine the agreement between the two-time measures of the same coder, we ran an intra-rater reliability test using Cohen’s Kappa (Kappa = 0.99).

Table S1. Population-level analysis (proportion of success). Correlations between fixed variables, mean human presence nearby, squirrel population and distance to the nearest footpath. (N = 15). Human presence nearby was the mean number of humans present within a 100 m radius around the puzzle box during each check. Squirrel population was the number of squirrels that participated in the experiment. Distance to the nearest footpath (m) was the shortest distance between the puzzle box and the nearest footpath. To avoid multicollinearity, variables with Pearson correlation r ≥ 0.5 (bold values) are not included in the same model during analyses.

| | Population | Distance to nearest footpath |
|---|---|---|
| Mean human presence nearby | 0.329 | |
| Distance to nearest footpath | 0.266 | -0.176 |

Table S2. Population-level analysis (proportion of success). Correlations between types of human activity (walking, dog walking, cycling, and playground activity) and distance to the nearest footpath (m) (N = 15). To avoid multicollinearity, variables with Pearson correlation r ≥ 0.5 (bold values) are not included in the same model during analyses.

| | Walking | Dog walking | Cycling | Playground activity |
|---|---|---|---|---|
| Dog walking | 0.35 | | | |
| Cycling | **0.94** | 0.31 | | |
| Playground activity | -0.20 | 0.01 | -0.09 | |
| Distance to the nearest footpath | -0.07 | -0.41 | -0.12 | -0.31 |

Table S3. Individual-level analysis (first success latency). Correlations between fixed variables, mean human presence nearby, squirrel population and distance to the nearest footpath. (N = 64). Human presence nearby was the mean number of humans present within a 100 m radius around the puzzle box during each check. Squirrel population was the number of squirrels that participated in the experiment. Distance to the nearest footpath (m) was the shortest distance between the puzzle box and the nearest footpath. To avoid multicollinearity, variables with Pearson correlation r ≥ 0.5 (bold values) are not included in the same model during analyses.

| | Population | Distance to nearest footpath |
|---|---|---|
| Mean human presence nearby | 0.39 | |
| Distance to nearest footpath | 0.22 | -0.13 |

Table S4. Individual-level analysis (first success latency). Correlations between types of human activity (walking, dog walking, cycling, and playground activity) and distance to the nearest footpath (m) (N = 64). To avoid multicollinearity, variables with Pearson correlation r ≥ 0.5 (bold values) are not included in the same model during analyses.

| | Walking | Dog Walking | Cycling | Playground activity |
|---|---|---|---|---|
| Dog Walking | 0.33 | | | |
| Cycling | **0.95** | 0.31 | | |
| Playground activity | -0.21 | 0.07 | -0.1 | |
| Distance to the nearest footpath | -0.02 | -0.43 | -0.08 | -0.33 |

Supplementary video

S1. An innovator, Panda, pushed a lever-end to solve the novel problem. This if it is close to a nut container, or pull (instead of push) the lever-end if it is far from the nut container so as to make a lever/nut drop (i.e. successful solving).
demonstrated the solutions for this problem are counter-intuitive to squirrels in which a squirrel (demonstrated by Mario here; also see video‘PST’ in the electronic supplementary material)

S2. A squirrel’s response to approaching humans while she was solving the task.
